# One Chromatin, Many Structures: From Ensemble Contact Maps to Single-Cell 3D Organization

**DOI:** 10.64898/2026.03.19.710883

**Authors:** M. A. Carignano, V. Backman, M. Kröger, Luay M. Almassalha, I. Szleifer

## Abstract

Understanding how chromatin folds in three dimensions remains challenging because most experimental assays capture low-dimensional projections of an underlying, highly heterogeneous polymer. Here we present an ensemble-based interpretive framework built on the previously introduced Self-Returning Excluded Volume (SR-EV) model, a minimal generator of nucleosome-resolution chromatin conformations based on stochastic return rules and excluded-volume geometry. Despite its simplicity, SR-EV reproduces key experimental signatures across scales: heterogeneous nanoscale packing domains resembling ChromEMT and ChromSTEM observations, sparse and highly variable single-configuration contact patterns analogous to single-cell chromosome conformation capture (Hi-C), and robust ensemble-level contact enrichment consistent with topologically associating domains (TADs). In this framework, Hi-C loop and TAD signatures are interpreted as ensemble-level statistical enrichments rather than invariant features of single-cell conformations. SR-EV is explicitly designed to generate large ensembles of complete three-dimensional chromatin configurations that can be projected consistently onto two-dimensional contact maps and one-dimensional genomic profiles. By introducing architectural-protein effects only through ensemble selection rather than explicit forces, SR-EV supports a separation between intrinsic polymer geometry and regulatory bias and suggests that TAD-like features can emerge as statistical enrichments rather than deterministic three-dimensional structures. Coordination number and probe-based accessibility computed directly from SR-EV provide a unified link between three-dimensional packing, two-dimensional contact maps, and one-dimensional genomic profiles. Together, these results establish SR-EV as a minimal and physically grounded reference framework for interpreting how heterogeneous chromatin ensembles give rise to multimodal experimental observables, while remaining consistent with the fact that chromatin organization is realized in individual cells.

**SIGNIFICANCE:** Chromatin domains, boundaries, and contact enrichments are often interpreted as fixed structural entities, even though most experimental measurements average over large and heterogeneous cell populations. The SR-EV framework shows that many of these features can be understood as emerging from minimal geometric rules combined with ensemble-level bias, without requiring explicit molecular interactions or deterministic folding mechanisms. By distinguishing single-configuration heterogeneity from ensemble-level statistical organization – including the emergence of packing domains– SR-EV supports an interpretation in which chromatin organization is realized in individual cells but must be analyzed through ensembles. This perspective clarifies the probabilistic nature of genome architecture and provides a tractable reference framework for interpreting multimodal genomic and imaging data.

## I. INTRODUCTION

Despite rapid experimental progress, achieving a mechanistic account of chromatin organization remains challenging because available modalities report different – and only partially overlapping– observables at different dimensionalities. Nanoscale electron-microscopy approaches (e.g., ChromEMT or ChromSTEM tomography) visualize physical packing within thin nuclear volumes at a few-nanometer resolution, revealing heterogeneous *packing domains*: irregular, conformationally defined clusters of nucleosomes embedded in more dilute regions, distinct from sequence-defined or topological domains, but carrying neither sequence nor topological information [1, 2]. ChromSTEM analyses indicate a continuous spread of domain radii (roughly 10–200 nm) and packing efficiencies, with local concentration profiles that grade smoothly from dense interiors to dilute surroundings and approach a volume fraction of ∼ 20% at the domain periphery [3]. Notably, the Self-Returning Excluded Volume (SR-EV) model produces a broad, non-bimodal spectrum of packing-domain sizes and local compaction levels under minimal geometric rules, offering a physically grounded baseline consistent with these imaging observations [4]. These measurements highlight substantial cell-to-cell variability at the single-configuration level and emphasize continuous heterogeneity in domain size and compaction rather than discrete ‘open’ and ‘closed’ states, motivating theoretical frameworks that can accommodate broad, non-bimodal packing distributions.

Chromosome conformation capture (3C) and its genome-wide extension, Hi-C, quantify pairwise proximity frequencies ordered by genomic coordinate rather than true spatial geometry. Population Hi-C reports ensemble-averaged contact frequency maps across many cells [5, 6], whereas single-cell Hi-C captures the specific set of contacts present in an individual nucleus [7, 8]. A contact map is typically defined as

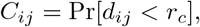

where *d*_*ij*_ is the spatial distance between loci *i* and *j* and *r*_*c*_ is a distance threshold. In population Hi-C, *C*_*ij*_ represents an ensemble probability derived from many cells, whereas in single-cell Hi-C it reduces to a binary record of whether a particular pair was captured as a contact in that nucleus. Because the mapping from any 3D conformation **X** to a contact matrix is many-to-one, contact maps cannot be inverted to a unique structure; ensemble averaging further collapses diverse single-cell geometries into a single contact-frequency matrix. Thus, while Hi-C is a powerful descriptor of connectivity, it offers only an indirect projection of spatial organization that must be interpreted statistically [9, 10]. Complementary imaging methods, such as multiplexed fluorescence *in situ* hybridization (FISH), provide direct spatial coordinates for selected genomic loci and therefore access true geometric distances, though at lower genomic coverage and throughput [11]. Together, these methods provide powerful but inherently reduced-dimensional views of chromatin that sample different aspects of single-cell structure [12].

Architectural proteins modulate chromatin folding by biasing probabilities rather than prescribing deterministic structures, thereby introducing variability across cells. Although sequence motifs bias where factors such as CTCF tend to bind, their actual occupancy fluctuates stochastically among nuclei. As a result, the effective positions of loop anchors vary from cell to cell, contributing to pronounced structural heterogeneity. Loop extrusion by cohesin, halted at convergently oriented CTCF binding sites, produces topologically associating domain (TAD)–like enrichment in ensemble Hi-C maps [13, 14]. However, live-cell imaging [15] and single-cell Hi-C [7, 8] show that such loops are transient and differently positioned across nuclei, indicating that individual cells realize only a subset of the configurations contributing to ensemble-level patterns. Consistent with this view, single-nucleus Hi-C measurements in *Drosophila* reveal substantial stochasticity in long-range contacts and demonstrate that TAD-like patterns emerge statistically despite pronounced cell-to-cell variability in individual genome folds [16]. Moreover, polymer models constrained by one-dimensional binding-site information reproduce population contact data only at the level of ensemble averages while generating a broad distribution of distinct single-cell conformations, as validated by Capture-C and FISH measurements [17]. These results underscore that agreement with population-level contact maps does not imply the existence of a unique or invariant three-dimensional structure. Further supporting an ensemble-based interpretation, CTCF depletion weakens/decouples contact-domain insulation [18] yet nanoscale packing domains persist [19], suggesting that intrinsic polymer dynamics, reversible protein-mediated interactions, and local crowding contribute substantially to domain formation [20, 21]. Ensemble features such as TADs therefore reflect statistically enriched configurations rather than invariant structures present in every cell [22]. Consistent with this interpretation, one-dimensional biochemical profiling (e.g., ChIP-seq) shows that architectural proteins and histone marks occupy non-random genomic positions that bias folding probabilities across populations without specifying unique three-dimensional conformations [23]. Importantly, such heterogeneity is not restricted to cell-to-cell differences: haplotype-resolved Genome Architecture Mapping (GAM) measurements reveal pronounced folding variability between homologous chromosomes within the same nucleus, reinforcing that chromatin architecture is inherently degenerate even under fixed cellular conditions [24].

Computational modeling provides an essential route to mechanistic insight, yet existing approaches capture only parts of this landscape [25]. Atomistic and mesoscopic simulations resolve nucleosome-level interactions with high fidelity but are restricted to fibers of at most tens to hundreds of nucleosomes [26–30]. Coarse-grained polymer models, including those implementing loop extrusion, operate on genomic segments ranging from hundreds of kilobases to tens of megabases –typically at a few kilobases per monomer– and can reproduce TAD-like enrichment in ensemble Hi-C maps [14, 31]. However, such models often prescribe specific folding mechanisms or rely on tuned effective interactions [32], which can reduce physical interpretability and dampen the stochastic variability evident in single-cell data. Bridging nanoscale packing, ensemble-averaged contacts, and single-cell variability therefore requires a model that is physically grounded, conceptually minimal, computationally scalable, and explicitly ensemble-aware.

To address these challenges, we previously introduced the SR-EV model, a minimal generative framework that treats the nucleosome as the basic unit and produces chromatin conformations using only self-returning steps and excluded-volume constraints. Without invoking attractive potentials, fitted interactions, or predefined looping mechanisms, SR-EV naturally yields broad, continuous distributions [4] of local packing density and packing-domain sizes –closely resembling ChromEMT and ChromSTEM measurements [1, 2]. The model is computationally efficient at megabase and chromosome scales, enabling direct comparison between single-cell–like heterogeneity and ensemble-level observables. Because SR-EV contains no intrinsic architectural proteins or sequence-specific cues, its unbiased ensembles are effectively randomized with respect to genomic locations of factors such as CTCF or cohesin. To incorporate such regulators without altering the model’s core physics, we introduce them as external constraints that select the subset of SR-EV configurations consistent with prescribed binding sites or loop-anchor geometries. This post-selection approach preserves the intrinsic packing behavior and stochastic variability of SR-EV while allowing protein-dependent structural enrichments –such as extrusion-stabilized loops or preferred boundaries– to emerge through conditional sampling rather than explicitly encoded forces.

This modularity enables simulations both with and without regulatory factors, helping to disentangle intrinsic polymer dynamics from protein-imposed organization and to capture variability between ensemble-averaged measurements and single-cell structures. In addition to reproducing qualitative features, SR-EV produces quantitative predictions for experimentally accessible observables –including contact-probability scaling, domain-size distributions, local volume fraction, and probe-based accessibility. These outputs enable comparison with population-level measurements such as Hi-C and electron-microscopy imaging, as well as with coarse-grained biochemical trends (e.g., ChIP-seq enrichment patterns), while maintaining nucleosome-level resolution [1, 2, 5, 6, 33]. Through this integrative structure, SR-EV provides a physically grounded framework linking molecular-scale interactions to chromosome-scale organization.

In this work, we use SR-EV to connect three scales of chromatin organization: nanoscale packing, single-cell structural variability, and ensemble-averaged genomic signatures. Slab-wise projections of single SR-EV configurations reproduce the heterogeneous packing domains observed by ChromEMT and ChromSTEM. Ensembles incorporating loop-anchor constraints yield TAD-like contact enrichment after averaging, even though individual structures remain diverse and sparse, mirroring single-cell Hi-C. Coordination number (CN) provides a direct link between three-dimensional density variation and one-dimensional compaction or accessibility profiles. Together, these features position SR-EV as a physically grounded framework for interpreting how heterogeneous 3D structures give rise to ensemble-level genomic patterns across experimental modalities.

## II. SR-EV MODEL

The Self-Returning Excluded Volume (SR-EV) model [4] describes chromatin as a polymer chain of nucleosomes grown through a stochastic process governed by only two elementary moves: *returns* to previously visited backbone positions and *jumps* to newly explored spatial locations [34]. At each growth step, the chain chooses to (i) perform a *Return* with probability

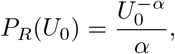

where *U*_0_ is the length (in units of 10 nm) of the previous backbone step and *α* > 1 is the folding parameter that controls the return–jump statistics; or (ii) perform a *Jump*, placing the next nucleosome at a distance *U*_1_ ≤ 20 drawn from

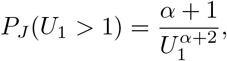

with an isotropically chosen direction. The chain is grown sequentially from a seed nucleosome under a spherical global cutoff of radius *R*, such that the occupied volume and the number of nucleosomes together achieve the target volume fraction *ϕ*. A short athermal relaxation with purely repulsive interactions removes any remaining overlaps. Together, the return and jump statistics, combined with the volume-fraction constraint, control the balance between extended exploration and compact self-returning, thereby shaping the distribution of packing-domain sizes. A full description of the model formulation, numerical implementation, and parameter exploration is provided in our earlier eLife work [4].

A notable feature of SR-EV is that its minimal rules – stochastic returns and excluded-volume constraints– are sufficient to generate heterogeneous three-dimensional structures containing compact packing domains interspersed with more open regions. These emergent domains arise at the level of individual configurations without any architectural proteins, attraction terms, or sequence-dependent inputs, reflecting purely geometric consequences of the growth rules. External regulators such as CTCF or cohesin are incorporated only after unbiased growth, through post-selection of configurations that satisfy prescribed anchor or binding geometries. This separation between intrinsic folding dynamics and extrinsic constraints preserves the broad structural variability generated by SR-EV while enabling controlled ensemble biases. Together, these properties make SR-EV a physically grounded baseline for exploring how minimal polymer rules give rise to packing domains, structural heterogeneity, and the ensemble signatures examined below.

## III. RESULTS

### A. Spatial heterogeneity and slab-wise packing analysis

To illustrate the internal organization produced by SR-EV, Fig. 1 shows a single chromatin configuration dissected into a series of adjacent slabs. Each slab is 50 nm thick and spans the full extent of the simulated volume in the other two dimensions, generating quasi–two-dimensional projections directly comparable to ChromEMT or ChromSTEM sections. Nucleosomes are colored by their coordination number (CN), defined as the number of neighbors within 11.5 nm, slightly above the ∼10 nm nucleosome diameter. CN values range from 0 in extended, dilute regions to 12 in locally compact domains, providing a direct visual readout of local packing density. Each slab reveals a heterogeneous mixture of packing domains spanning a broad range of sizes: large, high-CN aggregates resembling mature domains; intermediate-scale clusters; and small compact regions that may be interpreted as nascent or fragmenting domains.

**FIG. 1.**
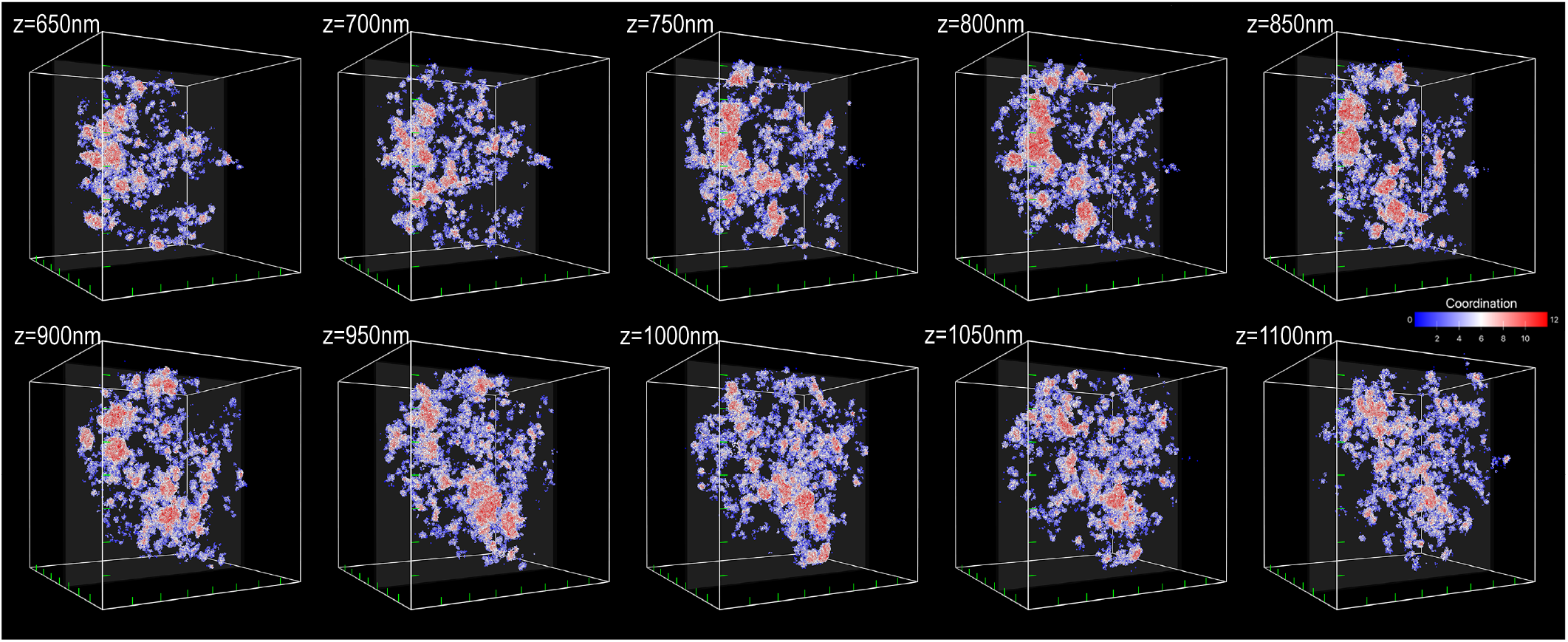
Spatial dissection of a single SR-EV chromatin configuration. A chromosome-scale SR-EV configuration consisting of *N* = 500,000 nucleosomes –comparable in size to human chromosome 15– was sliced into ten adjacent 50 nm slabs along the *z*-axis, spanning the full simulation volume in the other two dimensions. The configuration was generated with folding parameter *α* = 1.15 and volume fraction *ϕ* = 0.12. Nucleosomes are colored by their CN, defined as the number of neighboring nucleosomes within a cutoff distance of 11.5 nm, slightly larger than the ∼ 10 nm nucleosome diameter. CN values range from 0 (isolated nucleosomes) to 12 (densely packed local environments). Blue hues indicate loosely packed chromatin, red hues mark dense, compact domains, and white marks interfacial regions between these extremes. This slab-wise representation exposes the heterogeneous and hierarchical packing structure of a single chromatin configuration, in a format directly comparable to slab-based imaging techniques such as ChromEMT or ChromSTEM. Each slab contains a continuum of packing domains of varying sizes –large, highly compact regions, intermediate-scale clusters, and small, weakly formed domains– illustrating how SR-EV’s minimal rules generate the heterogeneous domain spectrum reported by ChromEMT and ChromSTEM. *This visualization illustrates how SR-EV’s minimal generative rules produce complex, domain-rich structures without imposed interactions*. See *Video S1* in the Supporting Material for a dynamic visualization.

This slab-wise representation makes the intrinsic heterogeneity of a single SR-EV configuration –our single-cell analogue– explicit: dense clusters coexist with more open regions, and interfaces between these structures give rise to intermediate CN values. Although CN is used here solely as a geometric packing metric, its spatial variation reflects the hierarchical structure emerging from SR-EV’s minimal rules and parallels the nonuniform packing observed experimentally. In addition, the slab-wise analysis exposes a spectrum of packing-domain sizes: large compact regions, medium-sized clusters, and small weakly formed domains coexist within the same configuration. Such graded, interdigitated domain populations mirror ChromEMT and ChromSTEM reports of continuous, non-bimodal distributions of local compaction rather than discrete architectural units. These slab-wise views define the heterogeneity present in any single SR-EV realization. The next step is to understand how much of this diversity persists –and how much is erased– when we shift from individual configurations to the ensemble representations used in population Hi-C.

### B. Loops and TADs

Individual SR-EV configurations are disordered 3D objects that lack a natural reference frame for direct structural averaging. Distinct realizations differ not only in local compaction and loop placement but also in global orientation and overall shape, such that any attempt to average configurations in real space is ill-defined. Even if configurations are aligned by their center of mass, straightforward averaging collapses structural information into a radially symmetric density profile, *ρ*(*r*), erasing domain boundaries and internal organization. As a result, the notion of an “average conformation” has no physical meaning for chromatin ensembles. Structural significance instead emerges only through averaging of reduced-dimensional observables –such as one-dimensional coordination number or accessibility profiles, and two-dimensional contact maps– that are invariant to global orientation. Accordingly, the analysis that follows is carried out in these projected spaces, with explicit connections to three-dimensional configuration space made only in specific examples. These quantities provide statistically robust descriptors of chromatin organization while faithfully reflecting the strong heterogeneity of individual configurations.

With this framework in mind, we next examine SR-EV ensembles of long chromatin segments generated under fixed loop constraints between specified genomic positions. The SR-EV model produces complete 3D configurations of chromatin chains (here *N* = 100,000 nucleosomes, corresponding to ∼ 18.5 Mbp), enabling direct inspection of individual realizations without imposing any additional alignment or ordering between configurations. In contrast to population Hi-C contact maps, which report pairwise contact frequencies averaged over thousands of nuclei and lack explicit geometric information, these full conformations make it possible to visualize internal variability in compaction, loop geometry, and domain-like organization at the single-configuration level. Despite the presence of fixed loop anchors, individual realizations remain highly heterogeneous in their spatial packing. Single configurations can display compact or extended conformations, one or multiple packing domains, or only weakly expressed internal structure, even when the same loop constraints are enforced. Thus, the existence of a loop in a given configuration does not uniquely define a packing domain or imply a well-defined higher-order organization at the single-cell level.

A widely discussed mechanism for TAD formation is the loop-extrusion model, in which cohesin complexes dynamically enlarge loops until arrested by convergent CTCF binding sites [14, 18, 35]. Such boundaries can stabilize loop positions and promote enriched contact domains at the population level. However, live-cell imaging and single-cell Hi-C measurements indicate that only a small number of extruded loops exist in any given nucleus and that their genomic positions vary substantially across cells [7, 15]. Consistent with a probabilistic interpretation, single-nucleus Hi-C studies show that individual genomes exhibit pronounced stochasticity in long-range contacts even when ensemble-averaged maps display reproducible TAD-like patterns [16]. These observations suggest that loop extrusion, when present, biases the statistical distribution of accessible folding states rather than uniquely determining a specific three-dimensional geometry in individual configurations. Uncertainty in the number, residence time, and spatial distribution of active cohesin complexes further limits fully deterministic descriptions of loop-mediated organization. Together, these considerations motivate modeling approaches in which loops are treated as statistical constraints whose effects emerge through ensemble averaging rather than as fixed geometric elements imposed at the single-cell level.

To introduce extrusion-like constraints into SR-EV *without modifying its intrinsic generative rules*, we generate large unbiased ensembles and subsequently post-select configurations containing loops anchored at prescribed genomic locations. This strategy mimics cohesin-mediated loop formation by enriching the ensemble for conformations consistent with specific CTCF–CTCF boundaries, while leaving the underlying stochastic growth process unchanged. Figure 2 illustrates this approach: four representative SR-EV configurations, each containing a 360 kbp loop (approximately 1,940 nucleosomes) extruded to completion between convergent CTCF sites, are shown. Despite sharing identical anchors and loop length, these configurations exhibit markedly different internal geometries. In particular, the same loop constraint can give rise to a single compact packing domain, several smaller domains, or a diffuse, low-density conformation across different realizations. This variability underscores that loop anchoring alone does not uniquely determine either the global fold or the internal domain architecture of the looped segment. Nucleosomes within the loop are colored by their coordination number (CN), and the mean CN decreases from left to right, illustrating that fixed-loop constraints still permit substantial variability in internal packing.

**FIG. 2.**
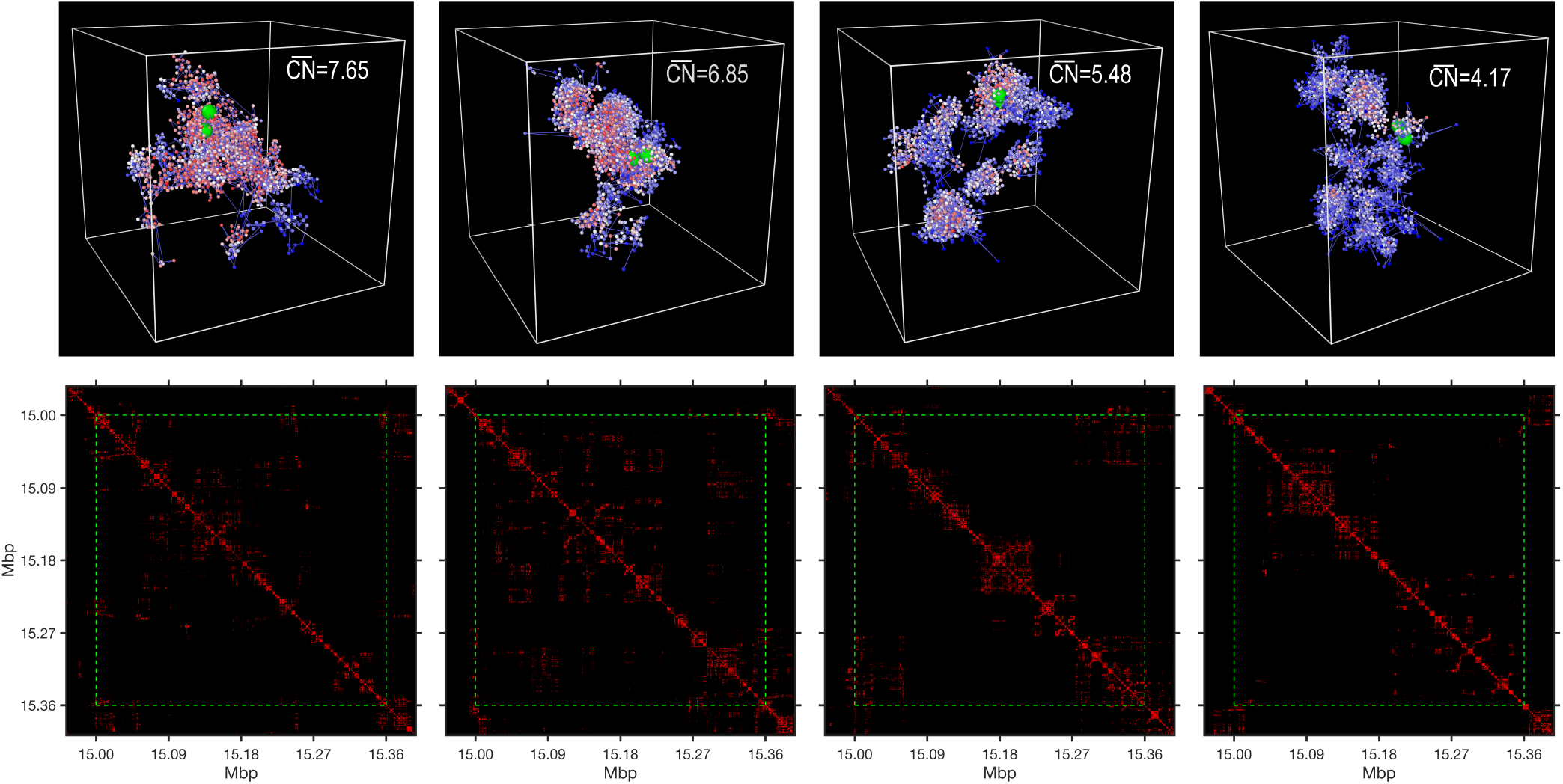
Variability of loop structures and contact patterns in SR-EV configurations. Segments of SR-EV chromatin configurations illustrating distinct structural outcomes of loop formation, along with their corresponding contact maps. **Top row:** 3D renderings of four *local 360 kbp segments* (spanning 15.00–15.36Mbp in the simulation index) extracted from full SR-EV configurations of *N* = 100,000 nucleosomes. Each segment contains a fully extruded loop anchored at fixed, convergently oriented CTCF binding sites (green spheres). Nucleosomes are colored by their coordination number (CN), reflecting local packing density. The mean value 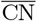 within the looped segment is reported for each configuration. Despite identical genomic anchors, the internal loop structure exhibits substantial variability across configurations, including differences in compaction, domain organization, and spatial arrangement. *Each looped segment also displays a heterogeneous mixture of packing domains –compact, intermediate, and dilute regions– highlighting that loop anchoring does not impose a unique internal domain architecture*. **Bottom row:** Contact maps computed from the same four looped segments. The green square highlights the genomic region between the loop anchors. These maps reveal the sparsity and heterogeneity of single-configuration contact patterns, showing that distinct 3D geometries can give rise to superficially similar contact signatures. Single-configuration contact maps are therefore not invertible to a unique 3D structure. Rotating views of the four looped segments are provided in *Videos S2–S5* in the Supporting Material.

These differences are not associated with changes in loop anchoring but instead reflect stochastic, configuration-dependent variations in local crowding and return statistics during chain growth. Importantly, the variability observed for this specific loop is not unique to this genomic position: analogous diversity is expected to arise for other cohesin-mediated loops along the same chromosome and across different genomic locations. As a consequence, an ensemble intended to represent a full chromosome would naturally comprise a complex superposition of many looped segments, each sampling a broad distribution of internal domain architectures. This hierarchical variability provides a plausible structural basis for the rich, multi-TAD contact patterns observed in population Hi-C maps.

Intermediate stages of loop extrusion can be modeled analogously. Figure 3 shows four configurations in which cohesin has extruded only part of the CTCF–CTCF span inside a fixed 360 kbp window, producing loops of 270, 216, 162, and 108 kbp (corresponding to roughly 1,450, 1,165, 875, and 585 nucleosomes). As before, green spheres mark CTCF anchors and yellow spheres mark cohesin positions. Although differences in overall compaction are modest in absolute terms, a clear trend emerges: shorter partially extruded loops yield lower average CN across the full anchor-to-anchor segment and exhibit sparser contact patterns. Importantly, the qualitative variability observed for fully extruded loops persists for these extrusion intermediates, with individual configurations again displaying compact, multi-domain, or diffuse packing within the looped region. These examples reinforce the structural and domain-level degeneracy inherent to loop-based mechanisms. What these examples reveal at the single-cell level, we now examine at the ensemble level: how do populations of structurally distinct looped states cooperate to produce robust TAD-like patterns?

**FIG. 3.**
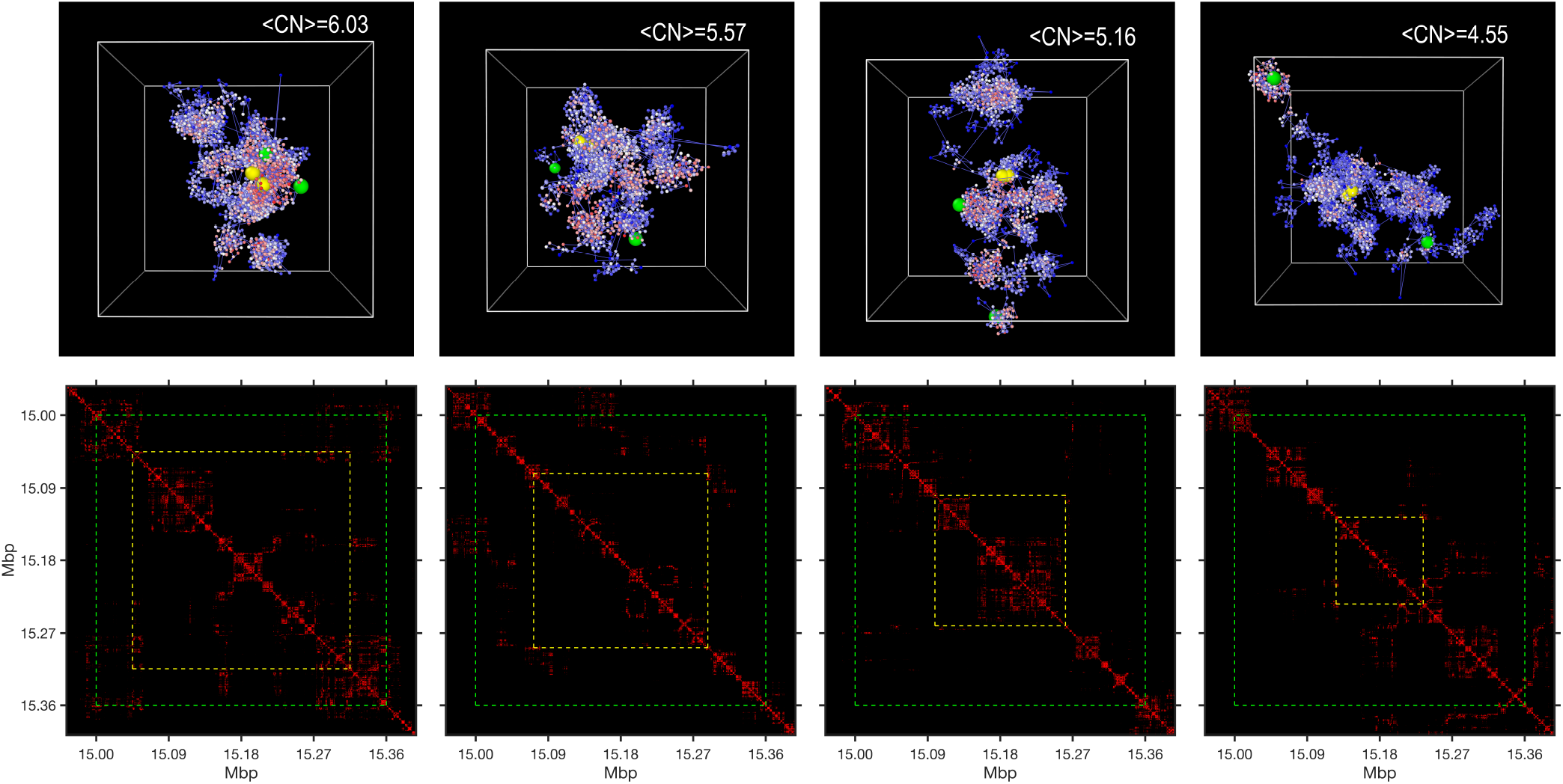
Intermediate stages of loop extrusion generate diverse 3D structures within the same genomic window. Four representative SR-EV chromatin segments illustrating partially extruded loops and their corresponding contact patterns. **Top row:** 3D renderings of four *local 360 kbp segments* (spanning 15.00–15.36Mbp in the simulation index) extracted from full SR-EV configurations of *N* = 100,000 nucleosomes. All segments contain a pair of fixed, convergently oriented CTCF binding sites (green spheres) defining the same 360 kbp domain, but cohesin has extruded only part of the span. From left to right, cohesin positions (yellow spheres) correspond to extruded lengths of 270, 216, 162, and 108 kbp, respectively. Nucleosomes are colored by their coordination number (CN), highlighting differences in local compaction across extrusion stages. Shorter extruded segments tend to yield lower 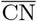 over the full 360 kbp window, illustrating that partial extrusion modulates compaction but still permits substantial structural variability. *Across extrusion stages, each segment exhibits a heterogeneous mixture of packing domains –from large compact regions to intermediate clusters and small, weakly formed domains– showing that loop growth influences compaction without imposing a unique domain morphology*. **Bottom row:** Contact maps computed from the same four partially extruded segments. The green square marks the full CTCF–CTCF span (15.00–15.36Mbp). Across the four examples, contact patterns remain sparse and heterogeneous –yet they exhibit similar coarse features despite the differing 3D geometries. These results reinforce that even within a fixed genomic window and fixed anchor positions, a broad range of single-cell structures can arise during loop extrusion. Rotating views of the four partially extruded segments are provided in *Videos S6–S9* in the Supporting Material.

To address this question, we generated SR-EV ensembles in which loops were required to form between prescribed genomic positions, mimicking convergent CTCF sites. In the ensemble shown in Figure 4, two adjacent domains were imposed: one loop of 720 kbp followed by a second of 360 kbp. Despite the broad range of three-dimensional geometries exhibited by individual configurations (Figures 2 and 3), the ensemble-averaged contact map displays a strikingly coherent pattern consisting of two well-defined TAD-like squares corresponding to these loop sizes. For comparison, an unbiased SR-EV ensemble sampled purely from intrinsic polymer rules without loop constraints produces smooth, largely featureless contact maps with no TAD-like enrichments (Supplementary Figure S1). The contrast between these two cases highlights a key principle: simple geometric constraints imposed on an intrinsically variable polymer ensemble are sufficient to generate strong, reproducible TAD-like patterns, even though no individual configuration resembles a canonical “TAD” in three dimensions. In other words, ensemble-level TADs reflect statistical enrichment at key nodes rather than uniquely defined volumetric structures.

**FIG. 4.**
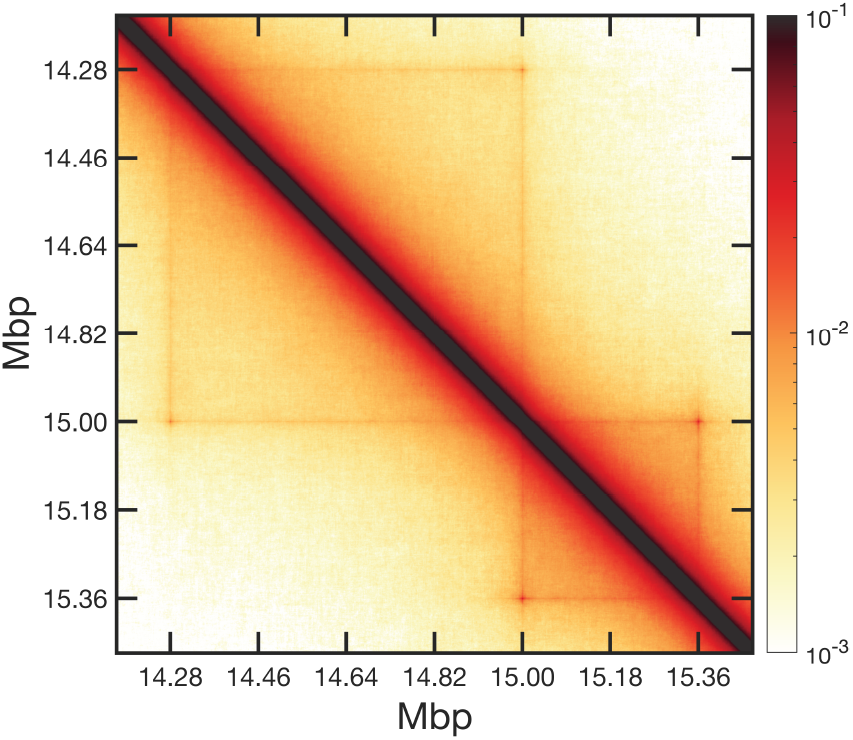
Ensemble-averaged contact map reveals two adjacent TAD-like domains. Contact-frequency map derived from 40,000 SR-EV configurations (*N* = 100,000) selected to contain either a 720 kbp or a 360 kbp loop anchored at fixed genomic positions. The resulting ensemble average displays two distinct square-like regions of enriched contacts –corresponding to the imposed 720 kbp and 360 kbp domains– while inter-domain interactions are comparatively depleted. Although individual configurations contributing to the ensemble remain sparse and structurally heterogeneous, their average yields robust TAD-like patterns that closely resemble population Hi-C maps.

### C. Linear Genomic Patterns

One-dimensional genomic profiles occupy a central role in chromatin biology. ChIP-seq maps histone modifications and architectural proteins; DNase-seq and ATAC-seq report local accessibility [36, 37]; and CUT&RUN or CUT&Tag provide high-resolution protein-binding landscapes [38, 39]. Although such measurements are often interpreted as indicators of specific structural features – such as loop boundaries, enhancers, or promoters– most datasets generated to date report signals that arise from population averages over highly heterogeneous three-dimensional chromatin conformations.

From this perspective, one-dimensional profiles represent a strong dimensional reduction: tens of thousands of distinct three-dimensional configurations are compressed into genomic traces. Most geometric degrees of freedom are lost in this projection—very different three-dimensional arrangements can contribute nearly identically to ensemble-averaged coordination number ⟨*CN* (*i*) ⟩ or accessibility ⟨*A*(*i*) ⟩ provided that their local packing statistics are similar. Individual SR-EV configurations, by contrast, exhibit sharp, irregular fluctuations that bear little resemblance to smooth ensemble-level trends. Consequently, one-dimensional genomic signatures should not be interpreted as literal features present in single cells, but rather as statistical projections shaped by the distribution of underlying three-dimensional states.

From a geometric standpoint, ensemble-level one-dimensional signatures are therefore naturally expected. If loop anchors bias an ensemble toward configurations in which, at the single-cell level, a genomic segment is spatially confined, then nucleosomes within that span must, on average, occupy a more crowded three-dimensional environment. Increased local crowding tends to raise the coordination number, while simultaneously tending to reduce the probability that a finite-sized probe can access nearby space.

Accordingly, ensemble-level contact enrichment should be accompanied by elevated ⟨*CN* (*i*) ⟩ and suppressed ⟨*A*(*i*) ⟩ within loop-delimited regions, with extrema expected at the anchors themselves. In this sense, correlated contact, coordination, and accessibility signatures arise as a direct geometric consequence of ensemble bias applied to a heterogeneous polymer, rather than as evidence for a unique or deterministic three-dimensional structure.

Figure 5 completes the link between ensemble-averaged contact maps and one-dimensional genomic readouts by projecting loop-constrained configurations onto linear genomic coordinates. The ensemble is constructed by combining two post-selected sub-ensembles: one consisting of configurations containing a 720 kbp loop and the other containing configurations with a 360 kbp loop, each imposed at fixed genomic positions. No individual configuration is post-selected to contain both loops simultaneously, although such configurations are not explicitly excluded. Using a total of 40,000 SR-EV configurations (each with *N* = 100,000 nucleosomes), we compute the ensemble-averaged coordination number ⟨*CN* (*i*) ⟩ and probe-based accessibility ⟨*A*(*i*) ⟩. In direct correspondence with the TAD-like contact enrichment, the one-dimensional profiles exhibit pronounced CN peaks at the respective loop-anchor positions and a modest elevation throughout each loop interior relative to the ensemble-averaged value of an unbiased SR-EV polymer grown at the same parameters.

**FIG. 5.**
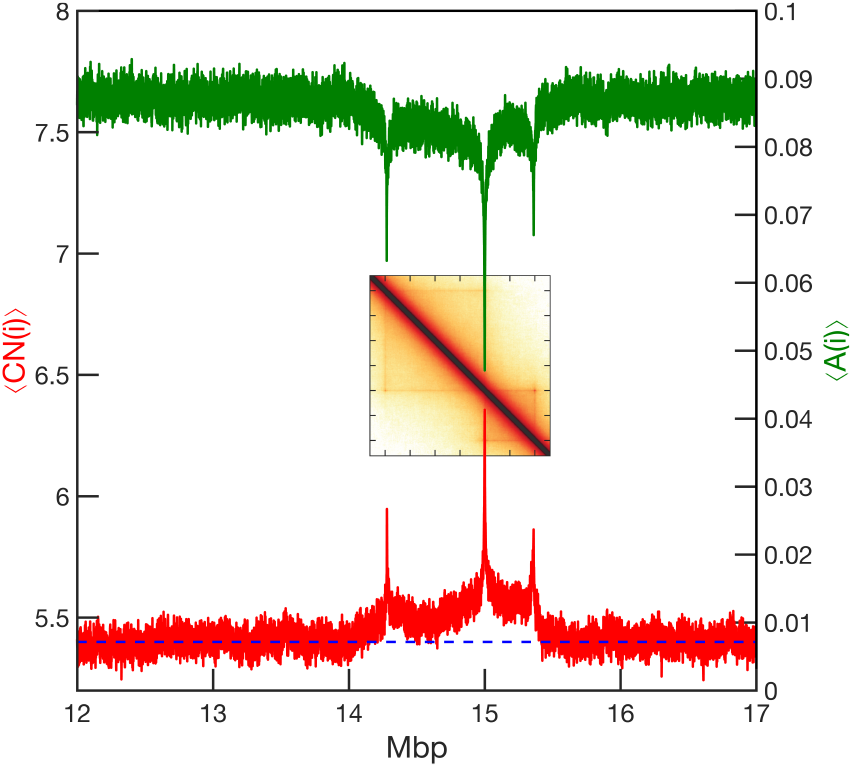
Ensemble CN and accessibility profiles across two loop-delimited domains. Ensemble-averaged coordination number (CN, red) and ATAC-like accessibility (green) computed from the same set of 40,000 SR-EV configurations (each with *N* = 100,000 nucleosomes; ∼18.5 Mbp) used to generate Fig. 4. Fixed loop anchors at 720 kbp and 360 kbp give rise to reproducible CN peaks at those genomic positions, marking the boundaries of the two loop-delimited domains. Within each domain, CN remains modestly elevated relative to the unconstrained baseline value of ∼5.4 (blue dashed line), reflecting persistent internal compaction across the ensemble. The accessibility profile shows the expected inverse relationship: regions of elevated CN correspond to reduced probe-based accessibility, with pronounced minima at the anchor sites and recovery to the baseline value of ∼ 0.095 outside the domains. Here, the unconstrained baseline ⟨*CN*⟩ ≈ 5.4 corresponds to the ensemble-averaged coordination number of an unbiased SR-EV polymer (i.e., without anchor-loop constraints) grown at the same volume fraction and parameter values, and serves as a reference for assessing compaction induced by loop constraints. *Inset:* Ensemble-averaged contact-frequency map (same data as Fig. 4) demonstrating that the 1D CN and accessibility signatures directly parallel the 2D contact enrichment imposed by the loop anchors.

The accessibility profile shows the complementary behavior: deep minima at the anchors, reduced accessibility across the loop interiors, and recovery to the baseline value of ∼0.095 outside the constrained regions. These correlated signatures reflect the repeated appearance of spatial confinement across the ensemble, which is amplified when projected onto one-dimensional genomic coordinates. This ensemble-level change in local packing is consistent with population observations that insulation and pronounced accessibility signatures (often peaks at CTCF/cohesin anchor sites) co-localize near loop anchors [6, 40].

Crucially, none of these smooth domain-level patterns are present in any individual SR-EV configuration. Single realizations instead display highly irregular, configuration-specific fluctuations in both coordination number and accessibility (Figures S2 and S3), underscoring that the linear signatures arise only through ensemble averaging. This observation suggests a broader question: if explicit loop constraints can generate extended domain-level profiles through ensemble bias, can similarly localized one-dimensional peaks—such as those observed in ATAC-seq, DNase-seq, or histone-mark data—also emerge through ensemble effects alone, without imposing loop constraints at the level of individual configurations? Experimentally, such peaks are often interpreted as evidence of stable enhancer–promoter contacts or precisely positioned architectural elements; here, we explore whether they may alternatively reflect conditional enrichment across a heterogeneous population of conformations.

To examine this possibility, we consider an unbiased SR-EV ensemble and construct a conditioned sub-ensemble based solely on a one-dimensional criterion. Specifically, we rank configurations by their probe-based accessibility within a narrow four-bead window (corresponding to ∼40–50 kbp) and retain only the top 20% of configurations. Each retained configuration remains a fully specified single three-dimensional conformation; however, no geometric constraints, loop anchors, or modified folding rules are introduced, and the underlying three-dimensional configurations are generated exactly as in the unbiased ensemble. The choice of the top 20% is illustrative rather than biologically calibrated, and serves only to demonstrate how ensemble conditioning can bias projected one-dimensional readouts under conditional sampling.

The resulting post-selected ensemble exhibits a sharp, localized peak in the ensemble-averaged accessibility profile ⟨*A*(*i*) ⟩ at the selected genomic position (Fig. 6), whereas the full ensemble shows a flat baseline of ⟨*A*(*i*) ⟩ ≈ 0.095. No single configuration contains the isolated full peak; rather, each contributes a partial and configuration-specific elevation. The pronounced one-dimensional feature emerges through statistical amplification across the conditioned ensemble. In a full genomic readout, multiple such peaks would correspond to a superposition of distinct conditioned sub-ensembles, whose relative weights may vary across cell types or cellular states.

**FIG. 6.**
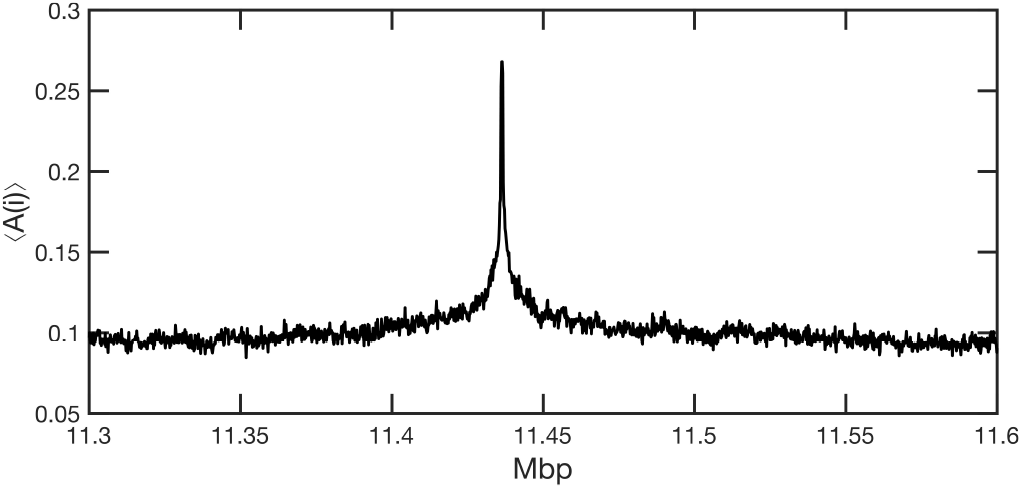
Localized accessibility peak generated by post-selection. Average ATAC-like accessibility ⟨*A*(*i*) ⟩ obtained from a post-selected SR-EV ensemble. An unbiased ensemble of 10,000 configurations (each with *N* = 100,000 nucleosomes) was ranked by probe-based accessibility within a narrow four-bead window (∼40–50 kbp), and only the top 20% (2,000 configurations) were retained. The resulting post-selected ensemble exhibits a sharp, isolated peak in ⟨*A*(*i*) ⟩ centered at ∼ 11.41 Mbp, whereas the full ensemble shows a flat baseline of ⟨*A*(*i*) ⟩ ≈ 0.095. No individual configuration contains the full peak; each contributes only a partial elevation, and the ensemble-average signature emerges solely through conditional sampling. This example illustrates how biologically recognizable one-dimensional patterns –resembling enhancer- or promoter-like accessibility features– can arise without geometric constraints or altered folding rules, reflecting the statistical amplification of heterogeneous three-dimensional configurations.

This example supports the idea that biologically recognizable one-dimensional signatures can arise through ensemble selection, without requiring unique underlying structures or explicit mechanistic remodeling. More broadly, it suggests that constructing an ensemble consistent with experimental observables is itself a nontrivial operation: different one-dimensional readouts may constrain overlapping but not identical subsets of three-dimensional configurations. In the Discussion below, we address how such constraints may interact, and how ensembles can be shaped to simultaneously satisfy multiple experimental projections of chromatin organization.

## DISCUSSION

A central message of this work is that chromatin organization is most naturally interpreted at the level of ensembles, rather than as a collection of reproducible three-dimensional structures. Experimental readouts of genome architecture are overwhelmingly low-dimensional –one-dimensional genomic profiles or population-averaged contact maps– yet they are often interpreted as reflecting underlying structural motifs. The results presented here support the view that such observables arise as statistical projections of a vast and heterogeneous space of three-dimensional configurations, even though chromatin function is realized within individual single-cell conformations. Consistent with this view, recent haplotype-resolved GAM experiments demonstrate extensive folding variability between homologous chromosomes within the same nucleus and cell type, indicating that multiple distinct three-dimensional conformations coexist under identical biological conditions [24].

A key clarification is that the framework explored here does not treat chromatin as a purely random polymer in vivo. Real chromatin exists in a molecularly rich environment and is continuously acted upon by architectural proteins, transcription factors, remodelers, and nuclear constraints. Rather, the model defines a minimal geometric and steric baseline: the space of conformations accessible to a nucleosome-resolved polymer under excluded volume and local self-returning growth. Within this interpretation, biological regulation may be viewed as biasing the statistical sampling of this pre-existing configurational space, selectively enriching subsets of single-cell conformations that satisfy functional constraints. In this view, ensemble formation reflects an active biological process: regulatory factors reshape how single-cell conformations are distributed across the ensemble, while the underlying geometric variability –and its parametric flexibility– remains intrinsic.

This ensemble-based viewpoint also clarifies the role of the post-selection examples explored in the Results. The 20% accessibility-based conditioning used to generate localized one-dimensional peaks is not intended as a biological claim about the fraction of cells, but rather as an illustrative example of a general mechanism. The results suggest that experimentally observed genomic features may be interpretable as arising when an ensemble is enriched for configurations that satisfy a given low-dimensional criterion. In a full genomic readout, different peaks would then correspond to a superposition of many such conditioned sub-ensembles, each contributing with a characteristic weight. These weights –and the resulting composite signal– are expected to depend on cell type, regulatory context, and physiological state; altered states, including disease, could plausibly correspond to systematic shifts in ensemble composition and/or effective parameters.

A central contribution of this work is to *apply and extend* the previously introduced SR-EV model as a practical platform for an ensemble-based interpretation of chromatin organization, while retaining access to complete single-cell three-dimensional conformations. In this role, SR-EV enables the generation of large ensembles of nucleosome-resolved three-dimensional conformations in which individual configurations remain fully interpretable, and that can be projected consistently onto orientation-invariant observables such as two-dimensional contact maps and one-dimensional genomic profiles. Each configuration represents a physically explicit single-cell realization, comparable to slab-based or section-based imaging readouts, while the ensemble formed by many such realizations supports direct comparison between heterogeneous single-cell structure and population-level signatures across experimental modalities. A further practical advantage is computational: the SR-EV model generates complete three-dimensional chromatin configurations at nucleosome resolution while remaining computationally efficient enough to sample large ensembles spanning megabase to chromosome-length scales. This tractability makes it feasible to quantify variability across many independent single-cell–like conformations, rather than relying on a small number of representative structures. This computational efficiency also makes it possible, in future extensions, to use SR-EV–generated chromatin ensembles as three-dimensional geometric substrates on which the spatial accessibility, encounter likelihoods, or effective action of regulatory proteins and enzymes could be assessed statistically, without explicitly modeling their molecular-scale dynamics. In this sense, SR-EV is not constructed to reproduce a single representative structure or a single readout, but to support interpretation of how experimentally observed 1D and 2D signals can emerge from biased sampling of a broad, heterogeneous configurational space. The examples presented here intentionally cover only small genomic segments and simple conditioning schemes; they are meant to demonstrate feasibility rather than completeness. The full complexity of chromatin organization –arising from many overlapping constraints acting across long genomic distances– can, in principle, be explored only within a framework that simultaneously resolves single-cell structure and ensemble-level statistics. By construction, different cell types or pathological states can be represented within the same framework through changes in ensemble weighting and/or parameter values, without altering the underlying generative rules.

This ensemble-based interpretation raises a natural question: can all experimentally observed onedimensional genomic features be simultaneously realized within a single chromatin conformation? The results presented here suggest that this is unlikely in general. Individual configurations display sharp, irregular, and configuration-specific fluctuations in coordination number and accessibility, reflecting local packing heterogeneity rather than a globally organized pattern. Accordingly, any single configuration is expected to contribute meaningfully to only a subset of the genomic features observed at the population level. Ensemble-averaged profiles therefore are most naturally interpreted as arising from the superposition of many partially overlapping single-cell contributions, rather than from repeated realization of a canonical or representative structure. Importantly, the framework presented here is not a dynamical description of chromatin motion. Instead, temporal variability observed experimentally –whether across cells or within the same cell over time– can be interpreted, in this scheme, as sampling across different single-cell configurations drawn from the same underlying ensemble.

From this perspective, biological organization is most naturally interpreted not in terms of a single structure, but in terms of the coverage statistics of an ensemble: how many configurations contribute to a given genomic feature, how frequently each configuration contributes, and how many distinct features a typical configuration supports. These quantities provide a compact way of summarizing how an astronomically large configurational space is sampled and constrained in practice, without requiring deterministic folding rules or uniquely defined geometries. In this view, chromatin organization is realized at the level of single-cell conformations, but becomes biologically actionable through ensemble statistics in a cellular population. Determining these coverage statistics can therefore be framed as an inverse problem: inferring the minimal ensemble composition consistent with a given set of low-dimensional experimental readouts.

Within the interpretive framework suggested here, the relationship between chromatin structure and regulation can be viewed through an ensemble lens. Architectural proteins, histone modifications, and transcriptional machinery need not be understood as imposing fixed three-dimensional geometries; instead, they can be interpreted as biasing the statistical distribution of accessible single-cell conformations by altering the relative weights of configurations within an ensemble. In this view, chromatin regulation operates through probabilistic sculpting of configurational space, rather than through precise geometric control at the level of individual structures. The framework developed here supports a separation between organizational features that arise from generic polymer geometry and those that reflect regulatory bias, while remaining agnostic about the specific molecular mechanisms by which such bias is implemented.

Taken together, these results suggest that understanding genome function may benefit from moving beyond the search for representative or canonical structures and toward quantifying how structural variability is distributed across ensembles. Key open questions concern how many configurations contribute appreciably to a given experimental signal, how many distinct signals a typical single-cell configuration can support, and how these statistics vary across cell types or regulatory states. Within the perspective developed here, such questions are naturally framed in terms of ensemble coverage and weighting rather than deterministic structure. This framing motivates the concluding section below, which summarizes the ensemble-aware interpretation while remaining consistent with pronounced single-cell heterogeneity.

## CONCLUSION

This study establishes an explicit separation between single-configuration structure and ensemble-level organization in chromatin. Individual realizations –our single-cell analogues– exhibit irregular, configuration-specific packing and contact patterns that do not generally correspond to canonical TADs or smooth genomic profiles. Yet, when configurations are analyzed through conditional selection or averaging across ensembles, statistically persistent features emerge. TADs, accessibility peaks, and boundary signatures therefore arise most naturally not as deterministic three-dimensional motifs, but as projections shaped by biased sampling of a heterogeneous configurational space.

This perspective naturally reframes the interpretation of experimental chromatin readouts. One-dimensional genomic profiles and population Hi-C maps are best understood as encoding not the structure of a typical cell, but information about how frequently different classes of configurations occur within a population. Regulatory mechanisms can then be interpreted as acting not by imposing fixed geometries, but by reshaping ensemble composition and coverage statistics. Changes in cell identity or disease state are thus most consistently interpreted as shifts in the weighting of underlying configurations, rather than as the emergence of entirely novel deterministic folds.

By providing a minimal and physically grounded framework in which three-dimensional packing and one-dimensional genomic signatures can be interpreted consistently, this work highlights the importance of ensemble awareness as a useful organizing principle for chromatin biology. More generally, it supports the view that biological function emerges from probabilistic organization across large configurational spaces, and that progress in the field may benefit from explicit consideration of ensemble composition, superposition, and variability rather than from reliance on representative structures alone.

Finally, this work frames chromatin organization as an inverse ensemble problem: given a set of low-dimensional experimental readouts, what distributions of three-dimensional configurations are consistent with them? Addressing this question benefits from models that are capable of generating large, diverse ensembles while maintaining internal geometric consistency across three-dimensional packing, contact maps, and genomic profiles. Although the examples presented here intentionally focus on limited genomic segments and simplified constraints, the underlying approach is in principle scalable to chromosome-scale systems and to combinations of multiple experimental modalities. Within such a framework, differences between cell types or disease states can be naturally interpreted as changes in ensemble composition and parameter regimes rather than as the appearance of qualitatively new structures. Establishing this ensemble-aware perspective provides a foundation for systematically integrating heterogeneous chromatin measurements and for quantifying how biological regulation reshapes genome organization through probabilistic, rather than deterministic, mechanisms.

## ACKNOWLEDGMENTS

The following grants have been supporting our work on this topic: National Institutes of Health grant R01CA228272 (IS, VB), National Institutes of Health grant U54 CA268084 (LMA, IS, VB), NIH training grant T32AI083216 and NIH career development award K23DK144661 (LMA), and Northwestern University Starzl Scholarship (LMA).

## Notes

https://drive.google.com/drive/folders/1w2rjMduSDyF6sbfgwurETEiIODYRN_K2?usp=share_link

### Competing Interest Statement

The authors have declared no competing interest.

