## Supporting Material for "One Chromatin, Many Structures: From Ensemble Contact Maps to Single-Cell 3D Organization"

### SUPPLEMENTARY VIDEO LEGENDS

Supplementary videos are available at Google Drive (Videos S1–S9).

#### Video S1. Dynamic slicing of a chromosome-scale SR-EV chromatin configuration

This video presents a moving 50 nm-thick slice through a three-dimensional SR-EV chromatin configuration consisting of 500,000 nucleosomes—approximately the size of human chromosome 15. The slice moves continuously along the $z$-axis, spanning the full simulation box in the other two dimensions. Nucleosomes are colored by their local coordination number (CN), defined as the number of neighboring nucleosomes within a cutoff distance of 11.5 nm. Blue regions correspond to loosely packed chromatin (low CN), red regions indicate densely compacted domains (high CN), and white regions mark the interface between these zones. This dynamic view mimics slab-based imaging techniques such as ChromEMT and ChromSTEM, while also revealing spatial variability along the depth of the nucleus—variability that cannot be captured by any single experimental slice.

#### Videos S2–S5. 3D rotation of SR-EV configurations with anchored chromatin loops

These four videos correspond to the top row of Figure 2 and show three-dimensional SR-EV chromatin configurations, each containing a 360 kbp loop anchored at fixed genomic positions (green spheres, representing CTCF sites). Nucleosomes are colored by their local coordination number (CN), which reflects local packing density. The internal structure of the loop region varies across configurations, highlighting the intrinsic variability of chromatin folding even under identical loop constraints. Each video presents a smooth rotation of the 3D conformation, providing a spatial perspective that complements the static snapshots shown in the figure. These dynamic views underscore the structural diversity of chromatin loops and the challenges in inferring full three-dimensional architecture from contact maps alone.

#### Videos S6–S9. Rotating views of SR-EV chromatin configurations with internal loops of varying length embedded within fixed-length segments

These videos correspond to the four configurations shown in the top row of Figure 3. Each configuration features a fixed 360 kbp genomic segment (marked by green spheres, representing convergent CTCF binding sites), within which an internal loop of 270, 216, 162, or 108 kbp is embedded. Yellow spheres indicate one possible position of cohesin along the loop extrusion axis. Nucleosomes are colored by their local coordination number (CN), reflecting local

### SUPPLEMENTARY FIGURES

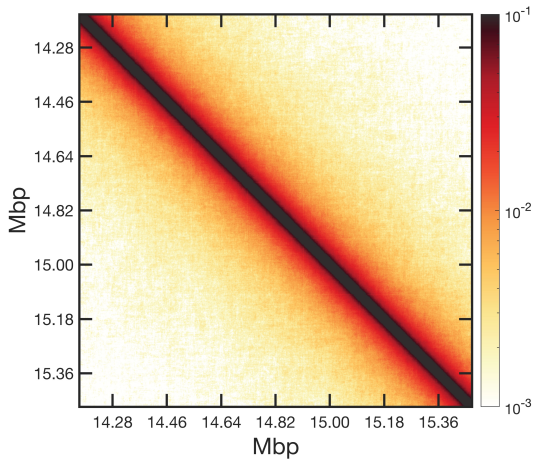


**Figure S1.** Ensemble-averaged contact-frequency map computed from an SR-EV ensemble generated without architectural-protein constraints or post-selection. Because configurations are sampled solely from intrinsic polymer geometry and excluded-volume rules, no preferred loop-anchor positions are imposed. The resulting map shows the expected smooth decay of contact probability with genomic distance and lacks square-like enrichments or boundary features, confirming that TAD-like patterns do not arise in SR-EV unless loop-anchoring constraints are applied. Color scale indicates contact probability on a logarithmic scale.

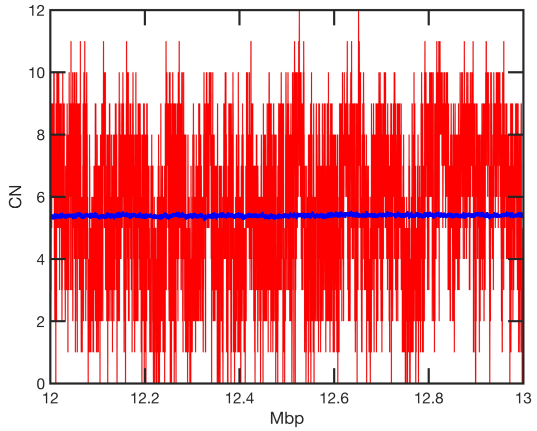


**Figure S2.** Coordination number (CN) profiles from SR-EV chromatin configurations. The red curve shows the CN$(i)$ trace from a single configuration, exhibiting strong bead-to-bead variability and intermittent dense regions. The blue curve shows the ensemble-averaged coordination number $\langle \mathrm{CN}(i) \rangle$, computed from the full SR-EV ensemble. The ensemble profile is smooth and reflects statistically persistent compaction patterns that emerge only after averaging thousands of heterogeneous conformations. This comparison illustrates that domain-like features in the average do not correspond to fixed structures in individual configurations.

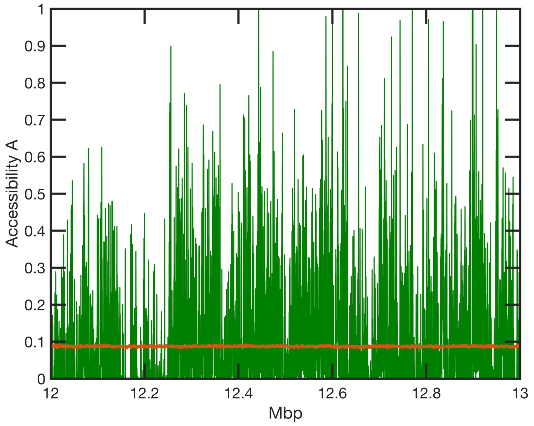


**Figure S3.** Accessibility profiles computed from SR-EV chromatin configurations. The green curve shows the single-configuration accessibility $A(i)$, which fluctuates strongly along the genomic coordinate and contains isolated low-accessibility pockets associated with transient dense regions. The orange curve shows the ensemble-averaged accessibility $\langle A(i) \rangle$, obtained by averaging over the full SR-EV ensemble. The ensemble trace is smooth and reflects statistically persistent patterns of compaction and openness that are not present in any individual conformation. This comparison highlights that accessibility-based genomic features arise only as ensemble-level statistical enrichments.
